# The shape of Cannabis sativa L. seeds for the differentiation of commercial medicinal cultivars from Argentina

**DOI:** 10.1101/2023.08.07.552192

**Authors:** Francisco Fernandez Torne, Tomas Bosco, Yanina L. Idaszkin, Gregorio Bigatti, Natahiel Garcés, Mariana Lozada, Rolando González-José, Federico Márquez

**Affiliations:** Institute of Biology of Marine Organisms (IBIOMAR, CONICET), Boulevard Brown 2915, U9120ACF Puerto Madryn, Chubut, Argentina; National University of Patagonia San Juan Bosco (UNPSJB), Blvd. Brown 3100, Puerto Madryn, Argentina; Patagonian Institute for the Study of Continental Ecosystems (IPEEC, CONICET), Boulevard Brown 2915, U9120ACF Puerto Madryn, Chubut, Argentina; Espiritu Santo University, Ecuador; Patagonian Institute of Social and Human Sciences (IPCSH, CONICET), Boulevard Brown 2915, U9120ACF Puerto Madryn, Chubut, Argentina

**Keywords:** Geometric morphometry, Chemotype, medical cannabis, INASE, CBD, THC

## Abstract

The Cannabis sativa L. plant has been used since ancient times as food, a source of fiber, and medicine, resulting in crosses that led to hybridization that currently does not allow for morphological differentiation among the three varieties of the genus (sativa, indica, and ruderalis). Currently, three chemotypes are differentiated based on their cannabinoid content (THC/CBD). Starting in the year 2023, seeds of two contrasting chemotype medicinal cultivars developed by CONICET (National Council for Scientific and Technical Research) and registered with INASE (National Institute of Seeds) can be commercialized in Argentina. In a previous study, we reported a relationship between the shape of Cannabis seeds and the chemical fingerprint associated with the chemotype. The objective of this study is to morphometrically characterize the seeds of two cultivars of Cannabis sativa L. with contrasting chemotypes: type I (high THC) and type III (high CBD). For this purpose, 2D geometric morphometrics based on landmarks and semilandmarks were used, allowing for the independent study of shape and size variation. Seed size between cultivars was compared using univariate statistics of an unbiased size estimator. To understand the magnitude and direction of shape change and determine shape characters that maximize separation between cultivars, a multivariate statistical approach was employed. Seeds belonging to the Malvina cultivar (type I cultivar, THC only) had, on average, smaller size and a rounded shape, whereas seeds from the Pachamama cultivar (type III, high CBD content) had larger size and a tendency towards an elongated oval shape. The use of a discriminant function based on seed shape allowed for over 97% correct assignments between cultivars. Our results could be used for implementing seed shape as a quality and authenticity seal for registered cultivars in the Argentine Cannabis market.

## 1. INTRODUCTION

The Cannabis plant is a dioecious herbaceous plant that has been used since ancient times by humans as food, a source of fiber, and medicine (Andre et al. 2016). The history of this plant has been marked by human interactions due to its multiple uses and customs over millennia, with its domestication traced back to archaeological data found in Central Asia at least 10,000 years ago (Robert et al. 2016; Long et al. 2017). Due to its diverse uses, the domestication of the Cannabis plant has been a long and complex process that has led to the emergence of different varieties. Over generations, Cannabis genetics have been selected and crossed to obtain specific traits of interest. Because of the extensive number of crosses and hybridizations, it has become increasingly difficult to classify and categorize different Cannabis cultivars (Jorasch 2020; Farag and Kayser 2015). Presently, there exist hundreds, even thousands, of cultivars, and this process of developing new genetics has been amplified in recent years with the expansion of the Cannabis industry. Currently, the trend is to classify Cannabis varieties into chemotypes based on the chemical fingerprint associated with the presence of active secondary metabolites (cannabinoids and terpenes) (Hazekamp et al. 2016), rather than by their morphology (indica, ruderalis, sativa). Notable among the cannabinoids found in plants are delta-9-tetrahydrocannabinol (THC) and cannabidiol (CBD). The most well-known Cannabis chemotypes are those predominantly high in THC (Type I); however, other chemotypes (Type II, with similar THC:CBD concentrations; Type III, predominantly high in CBD) have high therapeutic value. This classification has practical implications, as not only cannabinoids but also terpenes exert synergistic therapeutic effects (Pacifico et al. 2008). Therefore, pharmacological characterization is of paramount importance when using this plant as medicine in preparations with a full spectrum, as different cultivars can express different metabolites or in different proportions. In Argentina, in 2023, six medicinal cultivars of Cannabis sativa L. were developed at the CCT-CONICET-CENPAT facilities as part of the Interdisciplinary Cannabis Program implemented by the said center. These medicinal cultivars were registered by CONICET with INASE and are currently marketed as seeds of two cultivars named “Pachamama” and “Malvina”. The development of these seeds was achieved within the regulatory framework of Argentina regarding the medicinal use of the Cannabis plant and its derivatives, as well as the development of the medicinal Cannabis and industrial hemp industry (Laws 27,350/2017 and 27,699/2020). The CONICET cultivars present specific contrasting characteristics. “Malvina” is a Type I cultivar, with non-detected CBD percentages and THC: 11.4%, while “Pachamama” is a Type III cultivar, with a CBD:THC ratio of 16:1; with CBD percentages of 16.1% and THC of 0.96%. Additionally, these cultivars present a characteristic terpene chemical fingerprint; Pachamama predominantly presents terpenes: (-)-alpha-Bisabolol 28.9; (-)-Guaiol 20.7%; beta-Caryophyllene 16.9%; Myrcene 13%, and other terpenes below 10% such as Eucalyptol, alpha-Humulene, alpha-Terpineol, Beta-Pinene, Alpha-Pinene, d-Limonene, Linalool, Fenchyl alcohol, Borneol. On the other hand, Malvina’s major terpenes are beta-Caryophyllene 51.9%; alpha-Humulene 13.7; (-)-alpha-Bisabolol 10.6%, and other terpenes below 10%, such as Ocimene, d-limonene, (1S)-(+)-3-Carene, Myrcene, Beta-Pinene, Alpha-Pinene (see Resolution 238/2023, https://www.boletinoficial.gob.ar/detalleAviso/primera/281521/20230223). Evidence exists of the relationship between chemical characteristics and the morphology of Cannabis plants, although this relationship is often dependent on other characteristics associated with the cultivar (Clarke and Merlin 2013). However, there are morphological studies that use different anatomical structures, such as seeds (Small 1975; 1976), leaves (Anderson 1980), and cotyledon asymmetry (Small and Antle 2007), to determine different Cannabis variants. Many times, studies based on classical morphological characters, such as size, color, smell, leaf contour type, presence or absence of pores, veining, etc., yield contradictory results (Emboden 1974; Small and Cronquist 1976). In the last two decades, there has been significant growth in the use of geometric morphometrics on biological systems of all kinds, but there are only two works on C. sativa L. using this tool. Geometric morphometrics is defined as a fusion of geometry and biology, as it studies the shape of a specific object based on geometric space and multivariate statistical analysis. This method allows for the exploration of shape and size variations of objects in great detail, enabling the visualization of change, both in magnitude and direction, unlike classical morphometric analysis, which is based on the comparison of linear distances taken on the objects under study. The greater power and versatility of geometric morphometrics is due to the preservation of geometric information throughout the statistical analyses (Zelditch et al. 2012). Vergara and colleagues (2021) analyzed the correspondence between genetics and phenotype (classical characters and leaf shape) of Cannabis hybrids, observing that there is no association between these traits. In a study by Márquez and colleagues (2022), the association between seed shape of different unregistered C. sativa L. cultivars in Argentina and their respective chemotypes was studied using two-dimensional geometric morphometrics and multivariate statistical analysis. These authors found that seeds from mothers with a high THC concentration (Type I chemotype) had rounded and larger shapes compared to seeds from varieties with high CBD (Type III), which had elongated and smaller shapes. On the other hand, seeds with the presence of both THC and CBD (Type II) presented intermediate shapes and sizes. These differences allowed for effective chemotype distinction based on seed characteristics, proposing this technique as a promising tool for characterization and differentiation (Márquez et al., 2022). Given the importance of seed shape as an early indicator of cultivar type and the need to generate baseline data that allow for some form of commercial control, the objective of this study is to characterize and evaluate the seed shape of two medicinal Cannabis cultivars commercialized in Argentina using geometric morphometrics.

## 2. MATERIALS AND METHODS

### 2.1. The sample

A total of 220 stabilized feminized seeds were used, belonging to two commercial cultivars. Of these, 129 belonged to the Malvina cultivar and 91 to Pachamama. The seeds were obtained through backcrosses of female clones using the sexual reversion method through the application of exogenous growth regulators (Hall et al. 2012).

### 2.2. Determination of the shape and size of the seeds

The seeds were photographed under a Carl Zeiss binocular magnifying glass equipped with the AxioVision Rel.4.5 software (©Carl Zeiss Imaging Solutions) oriented with the abscission zone upwards, with the suture of the wall of the seeds. the radicle to the left (Fig.1). To capture the shape of the seed, a configuration of 3 anatomical *landmarks* and 10 *semi-landmarks on the contour* was used (Fig. 1), based on the proposal of Márquez et al. (2022). The term *landmark* refers to an anatomical “fixed point” and defines a discrete biological structure, while a *semilandmark* is a special type of *landmark* used to study the variation in structure contours between *landmarks* (Zelditch et al. 2012). Digitization of the Cartesian coordinates in the seed images was performed using a series of computer programs called TPS (*Thin Plate Spline*). Three modules of this series were used: TpsUtil (Rohlf 2017a) transforms one set of .jpg files to another. tps, then this file is opened in the TpsDig2 module (Rohlf 2017b) where the configuration of *landmarks* and *semi-landmarks* of each of the seeds is scaled and digitized. Subsequently, the *semilandmarks* are slipped to achieve their homology by means of a mathematical algorithm (“ *sliding* ”, an iterative process) that minimizes the bending energy *of* the TPS function, using the TpsRelw module (Rohlf 2017c). Finally, a Procrustes analysis was performed, which removes the effects of rotation, translation, and scale to preserve the information in a pure form (Zelditch et al. 2012). *Centroid size (* CS) was calculated as the square root of the sum of the squared distances from a set of *landmarks* to the centroid (Zelditch et al. 2012) and used as *a proxy* for the size of the seed. The CS retains all the size information, and in the absence of allometry (association between shape variation and seed size) it is the only size estimator that is not correlated with shape.

**Figure 1.**
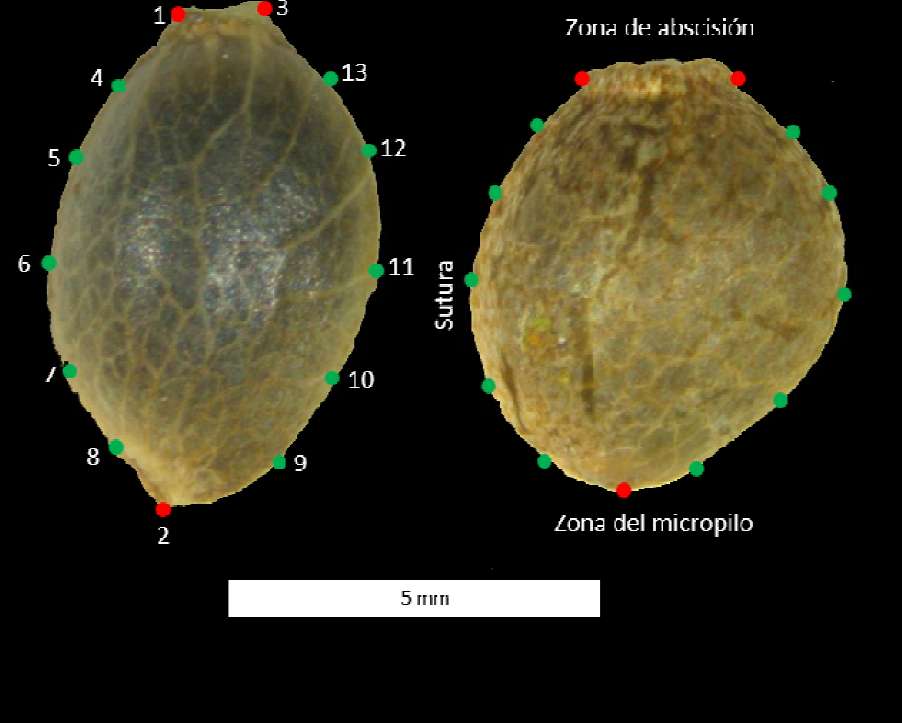
External view (perianth) of the longitudinal plane of the seed of the Pachamama (left) and Malvina (right) cultivar of *Cannabis sativa L. In both seeds, the configuration of landmarks* (red dots) and *semi-landmarks* (green dots) used can be observed. to capture the outline shape. *Landmarks* : (1) inflection point between the abscission zone and the suture of the fruit wall; (2) micropyle zone; (3) inflection point between zone of abscission and contour near the outer cotyledon; *Semilandmarks* : (4–8) semilandmarks placed equidistant between landmarks 1 and 2, in front of the suture contour; and (9-13) semi-reference points on the contour outside the cotyledon, between reference points 2 and 3. Scale bar=5 mm.

### 23. Statistical analysis of shape and size

Procrustes aligned coordinates and CS calculated in TpsRelw were exported to the MorphoJ software (Klingenberg 2011) to carry out multivariate statistical analyses. The presence of allometry was evaluated by calculating a multivariate regression between shape (aligned Procrustes coordinates, dependent variable) and size (CS, independent variable). To study the shape variation space (morpho-space) and determine the magnitude and direction of shape change, a principal component analysis was performed on the variance-covariance matrix of Procrustes distances. Then, in order to establish the components in such a way that they maximize the differences between the cultivar shapes and test the equality of mean shapes between them, a discriminant analysis and a Hotelling T 2 test were calculated. Finally, to evaluate the difference in means between the CS of the cultivars, a comparison of medians was made with a Kruskal-Wallis non-parametric analysis of variance test with the InfoStat software (Di Rienzo et al. 2011).

## 3. RESULTS AND DISCUSSION

### 3.1. Seed size

The frequency distributions associated with the size of the seeds of *C. sativa* L. for Malvina and Pachamama presented significant differences (Table 1). Pachamama seeds are on average larger than those belonging to the Malvina cultivar.

**Table 1:** Results of the Kruskal-Wallis test for the size comparison between the seeds of the Malvina and Pachamama cultivars.

| Variable | Cultivate | No. | Mean(mm) | OF | median(mm) | h | P-value |
| --- | --- | --- | --- | --- | --- | --- | --- |
| CS | Malvina | 129 | 5.78 | 0.44 | 5.84 | 50.23 | 0.0001 |
| CS | Pachamama | 91 | 6.31 | 0.5 | 6.37 |  |  |

This difference in size between cultivars is a useful characteristic for their identification. However, these results are in contrast to those found by Márquez et al. (2022), who found that the larger seeds belonged to a type I variety, while the smaller one registered was type III. Therefore, the size would be a reliable indicator for the separation between cultivars, but not for differentiating chemotypes.

### 3.2. Seed shape

Although the multivariate regression between shape and size (allometry) was significant, the percentage of association between cultivars was negligible (1.84%). This result was expected given that the seed is a structure that develops in a particular stage of the life cycle where, once mature, it does not present great growth variations. Similar results were found (absence of allometry) by Márquez et al. (2022) for another five varieties of *C. sativa* L. The first four principal components presented an accumulated variance of 90.19% of the total variation in shape. The shape variation explained by PC1, corresponding to 45.73% of the total variation, was associated with the slenderness of the seed. The positive extreme values presented more elongated shapes, with an expansion of the longitudinal plane of the seed, with a more projected apex and a restriction in the abscission zone. This shape space was occupied mainly by the Pachamama cultivar (chemotype III), while the opposite space (negative values) with rounded shapes, was occupied by Malvina seeds (chemotype I) (Fig. 2). The remaining principal component axes showed a high degree of overlap between cultivars.

**Figure 2:**
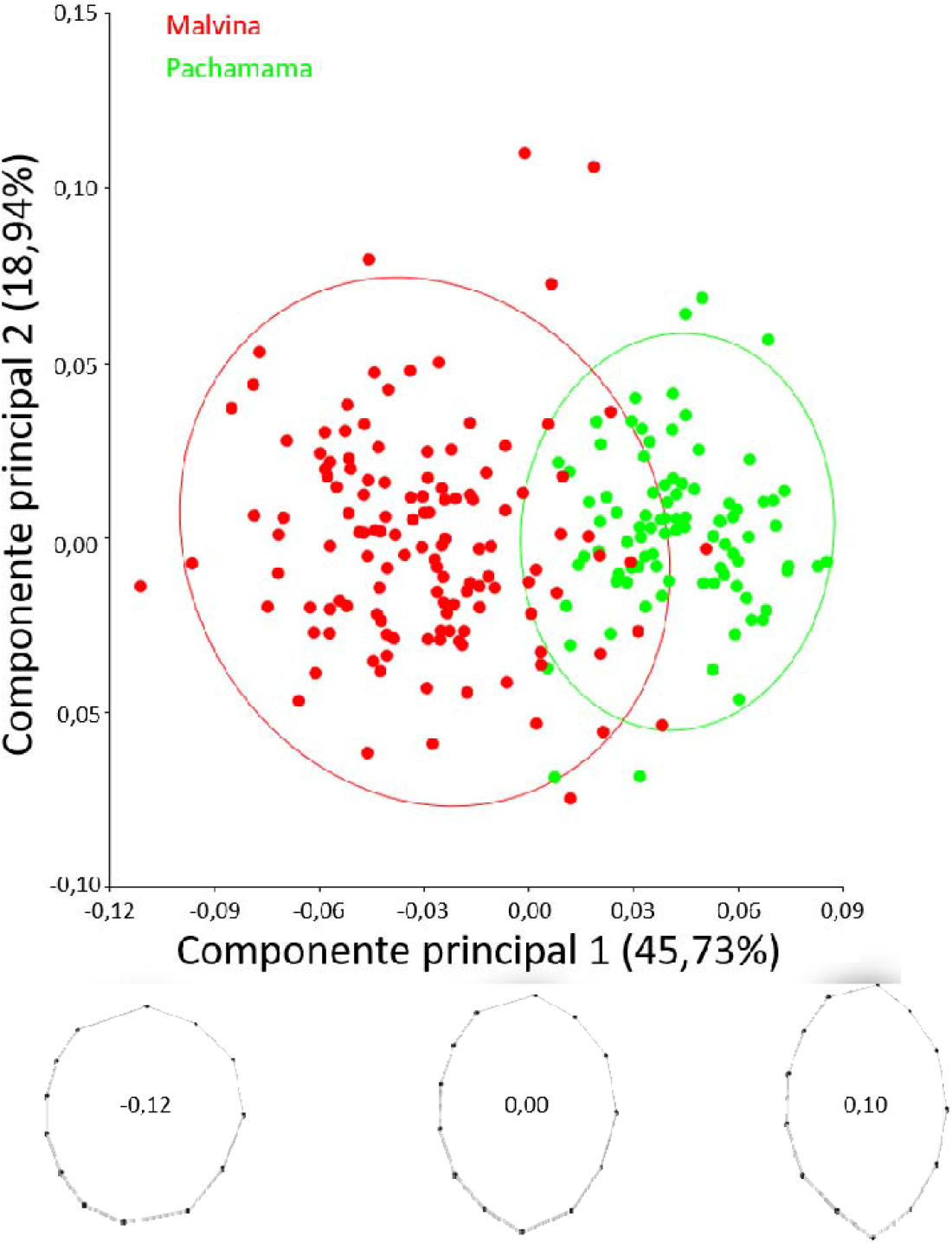
Analysis of principal components of the variations in the shape of the seed for 2 Argentine commercial cultivars of *Cannabis sativa* L. (Malvina and Pachamama). The ellipses correspond to the 95% confidence interval for the observations of each cultivar. The polygon plots show a positive value of -0.12, 0.0 (consensus), and 0.10 on the principal component 1 axis.

In agreement with the shape variations found throughout PC1, in the discriminant analysis (AD), the shape variation components that maximized the separation between the two cultivars were related to the slenderness/roundness of the seeds (Fig. 3). The comparison of the mean shape between Malvina and Pachamama showed a statistically significant difference (Table 2); (Hotelling T ^2^ : 1509.32, Mahalanobis distance: 5.31, p<0.0001). The cross-validated classification analysis showed that, for the two cultivars, 97.73 % of the seeds were classified correctly.

**Table 2:**
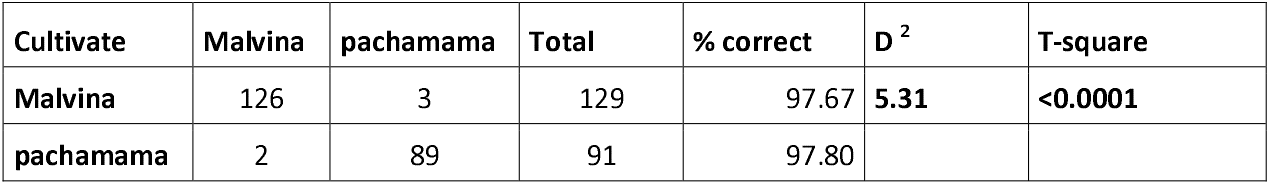
Cross-validated classification analysis showing the number of seeds assigned to each cultivar, using a discriminant function built from the shape of each cultivar. % Correct= percentages of assignments carried out correctly; D ^2^ = Mahalanobis distance.

**Figure 3:**
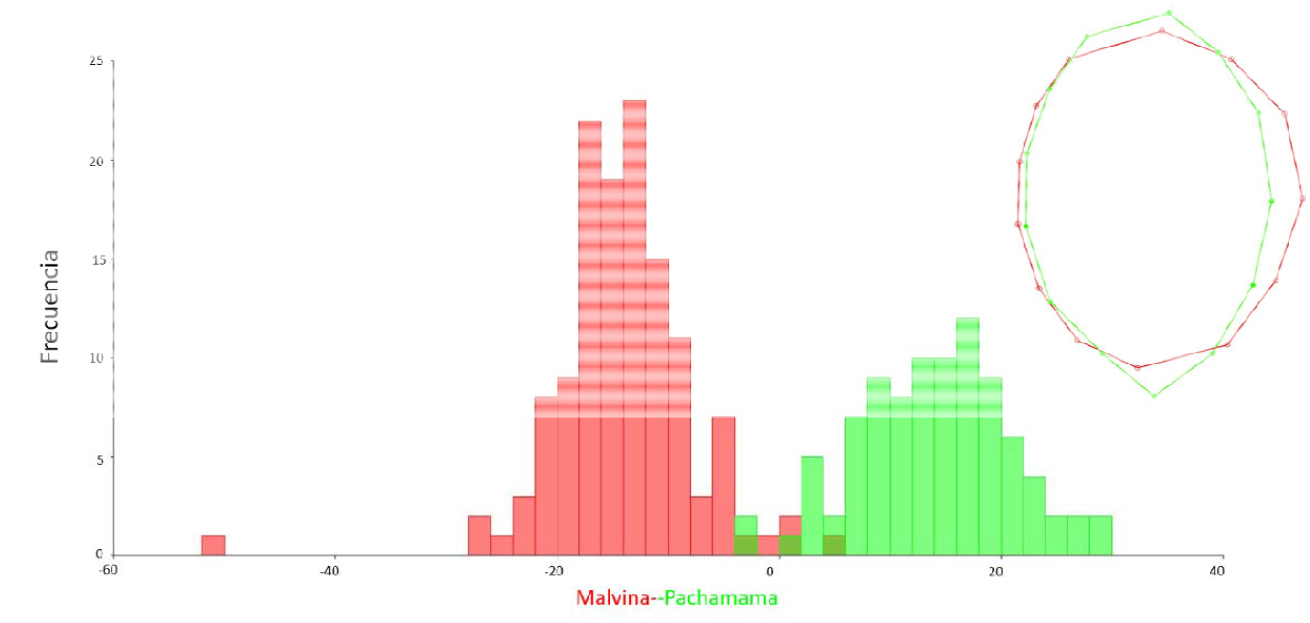
Diagram of frequencies predicted by the *jackknife* cross-validation test (leaving one out) between the seed form of the Argentine commercial cultivars of *Cannabis sativa* L. (Malvina and Pachamama). On the right, the graph of superimposed polygons represents the average form of both cultivars.

The main difficulties for a correct chemical identification of *C. sativa* L. cultivars are the cultivation time and the inputs necessary to obtain flowers, as well as the economic costs of chemically analyzing the different active components that constitute them. For this reason, being able to determine the cultivar and/or the chemotype early from the shape of the seed, through the use of geometric morphometry, is an alternative that could save time in the analysis and reduce costs, without losing effectiveness in the tests. assignments (Márquez et al. 2022). The results found in this work corroborate this hypothesis, since it was possible to determine with high percentages of effectiveness the assignments to the cultivars through the analysis of the shape of the seed. The commercial cultivar Malvina (type I-THC) presented rounded seed shapes similar to the type I varieties analyzed by Márquez et al. (2022), while the cultivar Pachamama (type III-CBD) showed elongated shapes, these being characteristics of the chemotype III varieties.

On the other hand, generating objective and quantified knowledge through a widely corroborated method such as geometric morphometry could allow its application in the control of specific characteristics intrinsic to each cultivar, for its registration in INASE. We believe that a discriminant function that nucleates all the seeds of the commercial cultivars registered by INASE in Argentina, could be used as a control for their recognition, against different stabilized varieties. Thus, the use of the shape of the seeds could be used to generate a seal of quality and authenticity of the cultivars registered in the *Cannabis market*.

## 4. ACKNOWLEDGMENTS

This work is part of the support scholarship within the framework of the Curriculum Improvement Program (ProMeC), of the province of Chubut (FFT). We thank the CD of the CCT CONICET-CENPAT, IBIOMAR and IPEEC for their support, as well as the personnel who collaborated with the Cannabis Interdisciplinary Program: administrative, maintenance, gardening, technological and electronic linkage personnel. This work was partially financed by the “Knowledge Nodes” project of the Ministry of Industry, Knowledge Economy and External Commercial Management and the Ministry of Productive Development of the Nation (RESOL-2021-1065-APN-SIECYGCE#MDP) and Project A6 of the Ministry of Science and Technology of the Nation.

## 5. CONTRIBUTION OF EACH AUTHOR

FFT: Conceptualization, data collection, analysis of results, writing of the original draft, writing, revision and editing. YLI: Conceptualization, resources, discussion of results, writing, revision and editing. TB: sample preparation, resources, writing, review and editing. GB: Sample preparation, resources, writing, review and editing. NG: Data collection, analysis of results. ML: Resources, writing, revision and edition. RG-J: Resources, writing, revision and edition. FM: Conceptualization, analysis of results, supervision, resources, writing, revision and edition.

## 7. FINANCING

This work was carried out within the framework of the Interdisciplinary Cannabis Program (https://cenpat.conicet.gov.ar/cannabismedicinal/), and with the partial financing of the project “Consolidation of the province of Chubut as a scientific and knowledge transfer node.” Cannabis-associated technology: comprehensive platform of services for the development of its productive chain” belonging to the Program “Nodes of the Knowledge Economy” (Ministry of Production, Argentine Republic), and project no. 6A of Cannabis Research and Development Projects (Ministry of Science, Technology and Innovation, Argentine Republic).

